# Hybrid MPL Scaffolds with Nanoscale Mechanobiology for Bone-on-Chip

**DOI:** 10.1101/2025.08.28.672808

**Authors:** Christoph Naderer, Eleni Priglinger, Martina Ramsauer, Cornelia Bergmayr, Tobias Gotterbarm, Dmitry Sivun, Jaroslaw Jacak

## Abstract

Engineering physiologically relevant 3D microenvironments is critical for studying cell behavior and advancing regenerative medicine. We present a new hybrid scaffold, fabricated via MultiPhoton Lithography (MPL), that integrates synthetic polymers (BisSR/CEA) with methacrylated collagen type I (Coll-MA) for single-cell enclosure and long-term culture. This is the first demonstration of a 3D MPL-printed biodegradable scaffold that mimics bone-like stiffness and allows spatially controlled, biodegradation-driven remodeling. The nanoscale feature size and mechanical properties are validated using Atomic Force Microscopy (AFM), while the nanoscale bioactivity of the scaffold is confirmed through Single-Molecule Localization Microscopy (SMLM). We track vinculin, a focal adhesion protein, with single-molecule resolution during mesenchymal stem cell (MSC) expansion and osteogenic differentiation. A new finding is time-dependent axial migration of vinculin clusters, independent of scaffold composition. Despite similar mechanosensing profiles, hybrid scaffolds significantly enhance osteogenic marker expression (collagen I, osteocalcin), revealing that scaffold bioactivity and geometry, not stiffness alone, direct stem cell fate. Cell expansion is highly dependent on scaffold composition, showing a biodegradation-driven remodeling over time. This platform offers a new tool to study cell-matrix interactions at the single-cell and single-molecule level and holds promise for Organ-on-Chip systems (*e*.*g*. bone-cartilage interface models), and personalized regenerative therapies.

## 1. Introduction

Mimicking and understanding tissue properties and functions is crucial in tissue engineering. Valuable approaches are Organ-on-a-Chip (OoC) systems, 3D-bioprinting, and spheroid/organoids cultures for investigation of cellular development *in vitro*.^[1–7]^ OoC systems incorporate microfluidic flow to simulate physiological conditions, including nutrient/waste exchange and mechanical stimulation. Cell bioprinting, on the other hand, enables precise spatial control over cell deposition to construct tissue-like architectures. However, OoC and bioprinting often lack the resolution to allow for single-cells analysis, limiting the ability to study cell-cell and cell-scaffold interactions at the micro-/nanoscale in a controlled 3D environment.

Recent developments in tissue engineering, particularly 3D-(bio)printed scaffolds, enable the studying and monitoring of cellular behavior in a complex environment, allowing researchers to assess the effect of the shape and topography of implants, or model diseases.^[8–15]^

The introduction of additive manufacturing has further advanced the field by enabling the fabrication of multi-material scaffolds.^[16]^ However, the integration of synthetic and protein-based materials for tissue engineering remains challenging. An advanced method in additive manufacturing is two-photon lithography introduced by Kawata in 1997 and has since been further advanced for high resolution 3D structuring of biomaterials.^[17]^ In two-/MultiPhoton Lithography (MPL), non-linear excitation of a photoinitiator triggers radical formation,^[18]^ leading to monomer crosslinking and solidification. MPL enables the fabrication of structures ranging from the nano-to macroscale,^[19–22]^ while achieving sub-100 nm lateral feature sizes,^[23–27]^ with a current axial resolution of ∼200 nm.^[28]^ In terms of 3D-(bio)printing, MPL outperforms technologies like stereolithography and electrospinning, commonly used in tissue engineering in terms of resolution or flexibility in geometry.^[29–34]^

For tissue engineering, MPL has mainly been used with synthetic materials.^[15,19,35–38]^ The biocompatibility of 3D synthetic scaffolds has been tuned by various strategies, such as the introduction of amino acid-based polymers,^[39,40]^ protein-adhesive photoresists and photochemical surface modification.^[10,41–43]^ The combination of nanostructuring and functionalization allows to observe cellular response on multi-material (*i*.*e*. hybrid) scaffolds.^[44]^ Recent developments implore protein-based photoresists to fabricate 3D scaffolds.^[45]^ Collagen, the most abundant structural protein in the ECM, due to its ability to support cell adhesion, migration and differentiation, is a particularly promising protein for 3D-(bio)printing,^[46]^ 2D guided cell growth,^[47–49]^ 3D patterning within hydrogels and for MPL fabrication.^[50,51]^

Next to geometrical factors, affecting cell growth and attachment,^[52–54]^ mechanical properties play a crucial role in influencing cell responses.^[55,56]^ Especially bone tissue presents a unique challenge due to its heterogeneous mechanical properties.^[57–59]^ Substrate stiffness can be adjusted by altering composition or applying external stimuli, a common strategy in research for studying changes in cell behavior.^[60–64]^ Mesenchymal stem cells (MSCs), stromal cells with multipotent differentiation potential *in vivo* and *in vitro*, are of high interest especially in tissue engineering, regenerative medicine and clinical applications.^[65–68]^ For example, hydrogel stiffness induces stem cell differentiation,^[69,70]^ increases reactive oxygen species expression in MSCs,^[71]^ and triggers phenotype changes in macrophages,^[72–74]^ all of which aid in tissue repair. While the influence of 2D substrate stiffness on stem cell fate is well established, recent studies have demonstrated that 3D microenvironmental cues - including matrix stiffness, viscoelasticity, geometry, and degradability—critically regulate mechanotransduction and lineage commitment.^[75–78]^

However, most prior studies have investigated bulk or multicellular 3D systems, limiting mechanistic insight at the single-cell level. A platform that enables truly 3D, dynamically remodeling scaffolds with nanoscale control of geometry and stiffness could allow direct quantification of single-cell mechanosensing. Thus, revealing how local scaffold architecture and bioactivity - rather than bulk material stiffness - govern mesenchymal stem cell osteogenic differentiation. Furthermore, leading towards establishing a new framework for studying cell–matrix interactions in physiologically relevant environments.

The mechantransduction allows cells to perceive matrix stiffness (*i*.*e*. vinculin-mitogen-activated protein kinase 1 (MAPK1) mediated focal adhesion spot activity), while guiding the commitment towards specific differentiation lineages.^[75,79,80]^ Vinculin, an actin filament-binding protein, regulates and reinforces integrin binding, links the actin cytoskeleton to the ExtraCellular Matrix (ECM), as well as being a key component in the mechanotransducting vinculin-talin complex.^[81–83]^

Studies show that stem cells spreading area and shape strongly influence osteogenic differentiation, with larger areas and lower circularity enhancing bone formation.^[84,85]^ While 2D micropatterning offers precise control, there is a need for 3D platforms that better mimic bone microenvironments. Such tissue scaffolds must balance structural support, mechanical properties, biodegradability, and biocompatibility while supporting cell growth.^[86]^ Current *in vitro* models often lack the hierarchical complexity of native bone, such as osteocyte lacunae or canaliculi, limiting their ability to capture fine-scale mechanosensing and differentiation processes.

Advances in single-cell technologies and *in vitro* models have enabled physiologically relevant stem cell studies,^[87]^ such as identifying tissue-specific MSC subpopulations influenced by ECM-associated heterogeneity.^[88]^ While biomimetic scaffolds are designed to enhance stem cell function and tissue regeneration,^[89]^ single-cell enclosure methods like microfluidic and encapsulation platforms, which control individual cell microenvironments, remain underexplored.

We report, for the first time, 3D hybrid scaffolds integrating collagen type I and biocompatible polymers, fabricated via MultiPhoton Lithography (MPL) with a biodegradation-driven, dynamically remodeling geometry inspired by bone. Unlike conventional 3D scaffolds or bioprinting approaches, which lack nanoscale resolution and dynamic remodeling, these scaffolds enable precise control over single-cell microenvironments. These scaffolds enable single-cell enclosure and spatial control, minimizing migration and allowing detailed study of cell-matrix interactions. Using AFM, SMLM, and confocal imaging, we characterized nanoscale mechanical properties and biofunctionality, while cluster-based analysis of vinculin distributions quantified 3D mechanosensing and focal adhesion organization at single-cell resolution.

We demonstrate that scaffold remodeling directs cell spreading, ECM deposition, and enhances osteogenic marker expression, revealing how geometry and bioactivity—rather than bulk stiffness alone—guide stem cell differentiation. MPL further enables tunable micro- and nanoscale stiffness, broadening future applications in mechanobiology and stem cell fate studies. The platform is also suitable for controlled 3D osteogenesis, highlighting its potential for bone-on-a-chip and Organ-on-Chip tissue engineering applications. This modular platform also allows future integration of more complex architectures, enabling detailed studies of cell-matrix interactions in physiologically relevant 3D microenvironments.

## 2. Results

### 2.1. 3D hybrid scaffold designed for single cell enclosure and observation

Imitating complex tissue properties is key to understanding cellular development *in vitro*. First, we focused on designing a 3D scaffold to support cell growth within a controlled 3D microenvironment, where material composition, geometry, and mechanical properties influence cell differentiation. The scaffold was specifically designed to enable the investigation of cell-matrix interactions at the single-cell level in this 3D context. We tailored a stadium-like structure (500x200x30 µm^3^, **Figure 1**a) inspired by bone with pore sizes supporting osteoblast mineralization and differentiation, and high mechanical stability using the commercial monomer Ormocomp®.^[19,90]^ The side walls of the stadium-like structure were designed with a tilt of 70° for an increased opening surface for easier cell seeding. The stadium-like structure segregates cells within the scaffold, minimizes direct cell-cell interactions, and prevents cell migration in and out of the scaffold (see **Figure S1**). The structure provided structural support for the more delicate microenvironments, which were printed in a second step inside of the stadium (SEM insets in Figure 1a, for stadium-like with cell cages see **Figure S2**). We printed symmetrical cell cages with dimensions of 45×45×24 µm^3^. The cell cages were fabricated 6 µm lower than the stadium-like structure to ensure cell segregation (stadium-like structure height: 30 µm, cell cage height: 24 µm). Hard cell cages were comprised of biocompatible synthetic polymer bars in the x- and y-direction, while hybrid cell cages were a combination of synthetic polymer bars in the x-direction and methacrylated collagen (Coll-MA) bars in the y-direction (Figure 1b).

**Figure 1.**
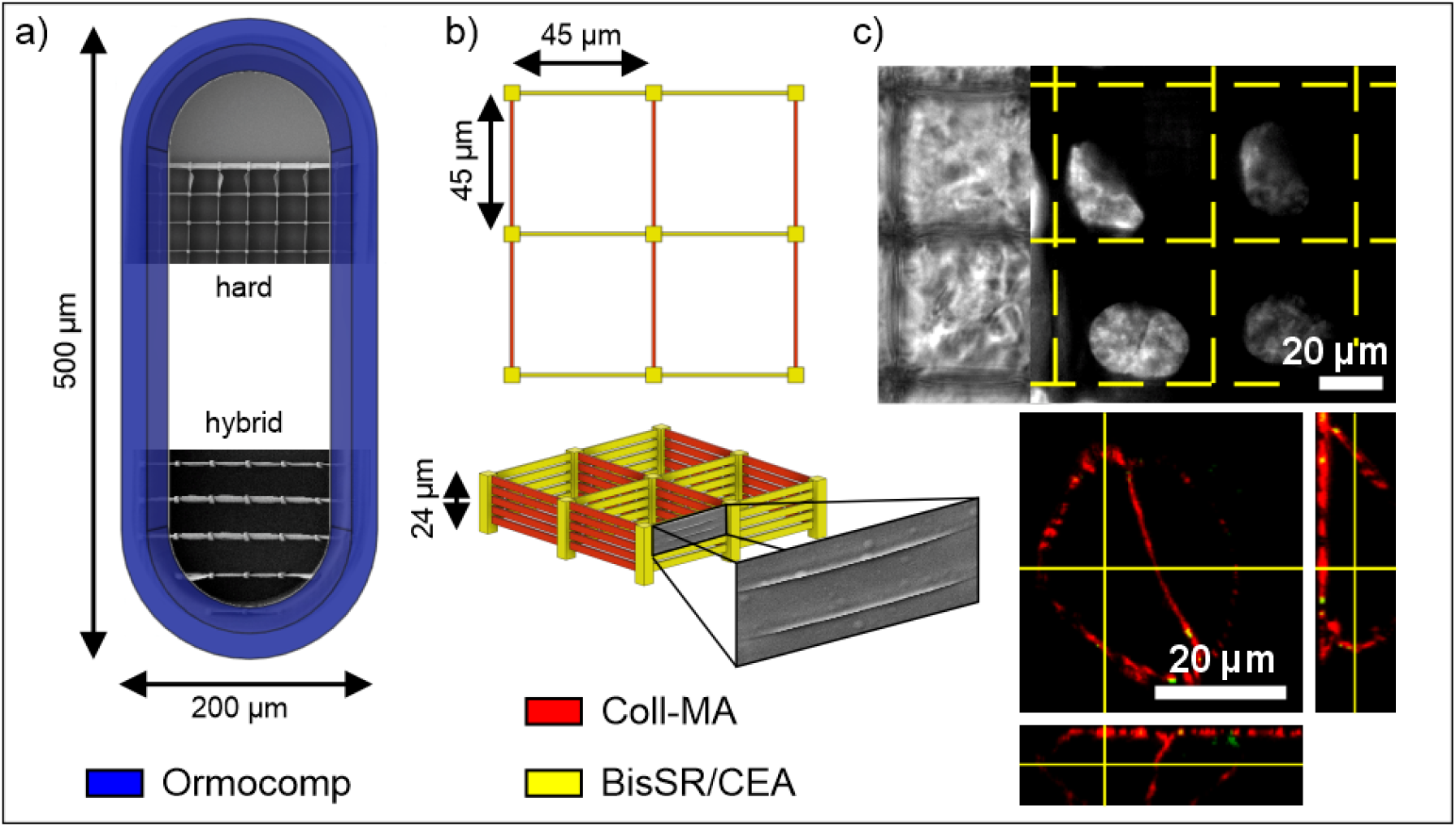
3D scaffold design for single-cell analysis. a) Schematic drawing of the stadium-like structure, designed to separate cells growing around the scaffolds from cells growing within the scaffolds. Insets show SEM images of hard cell cages (BisSR/CEA) and hybrid cell cages (BisSR/CEA and Coll-MA). The hard material BisSR/CEA supports the soft biofunctional methacrylated collagen type 1 bars (Coll-MA) incorporated in hybrid cell cages. b) Schematic drawing of hybrid cell cages composed of 5x5×24 µm^3^ pillars and 45x1x3 µm^3^ bars consisting of BisSR/CEA and Coll-MA, respectively. Axially written bars are separated by 1 µm and start 3 µm above the substrate to ensure cell-cell contacts between neighboring cells within the cell cages. The schematic 3D view of the cages with an additional inset highlights an SEM image of the bars separated by a gap. c) White light image (left) of MSC growing in cell cages 6 days post-seeding and fluorescence image of cell nuclei stained with DAPI. Single cells were housed within the scaffold. The lower confocal fluorescence image shows the actin cytoskeleton labeled with phalloidin Alexa647Plus (red) and vinculin labeled with anti-vinculin IgG Alexa488 (green). Schematics are not drawn to scale.

To facilitate cell-cell contact and communication (*i*.*e*. paracrine signaling), crucial for cell growth and development,^[91]^ the cell cages were designed with 1 µm gaps between cage bars with an additional 3 µm gap to the substrate. Furthermore, this ensures a genuine 3D architecture ensuring that cells interact with all surrounding walls, a feature not achievable with simpler planar designs. Combining biocompatible Bisphenol A-Glycidyl MethAcrylate (Bis-GMA), ethoxylated bisphenol A dimethacrylate (SR348C) (BisSR) and protein-binding 2-CarboxyEthyl Acrylate (CEA) in a weight ratio of 1:3.5:0.1 allows for coating of the hard cell cages with collagen prior to cell seeding.^[10,92]^ We utilized the biodegradability of Coll-MA to fabricate a 3D hybrid scaffolds with a progressively changing geometry to investigate cellular development. The cell cages provided the required support for cell attachment and expansion in 3D for 21 days of cell culture. The consistent enclosure of single cells within the scaffold was visualized by cell nuclei staining using 4’,6-DiAmidino-2-PhenylIndol (DAPI). The actin cytoskeleton was labeled with phalloidin Alexa647Plus as an indicator for cell morphology/spreading and cytoskeleton organization (red in the lower panel in Figure 1c). The focal adhesion-associated protein vinculin, involved in mechanotransduction, has been visualized using anti-vinculin IgG Alexa488 to assess cell-matrix adhesion (green in the lower panel in Figure 1c). We determined the best-suited grid size for single-cell spreading via visualization of actin cytoskeleton in tested grid sizes: 30 µm, 35 µm, and 45 µm (see **Figure S3**).^[53,56]^ Initially, 30 – 40 % of cell cages were filled during seeding. Cell cages with the 3 µm gap housed cells in all cages 6 days post-seeding (Figure 1c), while for cell cages with no gap to the substrate it took up to 10 days for the same occupancy. Additional exclusion of the 1 µm gaps between bars resulted in impeded cell expansion (see Figure S3, 30 µm grid size). Herin, we show a 3D hybrid scaffold allowing consistent cell enclosure and multi-color imaging at the single-molecule level, enabling monitoring of cellular spreading and cell-scaffold interactions at the single-cell level over time.

### 2.2. Functionality and mechanical properties of collagen-based structures

While the development of cells in 3D environments is a growing field of research,^[93–95]^ cell behavior within biodegradation-driven scaffolds remains poorly understood. To address this gap, we optimized next to the scaffold geometry, also the material composition, with a particular focus on mechanical properties and the biofunctionality of the structures. We quantified the mechanical properties of BisSR/CEA and Coll-MA-based structures and developed a strategy to match the variations of the Young’s modulus (*YM*) observed in bone and other tissues (ranging from kPa to GPa).^[57,75]^ To achieve this, we utilized Poly(Ethylene Glycol) DiAcrylate (PEGDA) as a crosslinking agent to precisely modulate elasticity.^[45]^ Firstly, we fabricated suspended lines, as previously outlined by Buchroithner *et. al*.^[19]^ to achieve 3D features minimizing surface effects, between Ormocomp® bars for mechanical characterization with AFM.

We used nanoindentation to characterize the *YM* of the components of the hybrid scaffold and to assess the modulation of *YM* of Coll-MA with the addition of PEGDA. As a reference, BisSR/CEA exhibited a *YM* of 81 ± 10 MPa (STD). With the addition of 5 wt% PEGDA to Coll-MA, the *YM* increased by a factor of ∼7 compared to pure Coll-MA lines (Coll-MA/PEGDA: *YM* = 368 ± 150 kPa, Coll-MA: *YM* = 49 ± 32 kPa) (**Figure 2**a). The high standard deviation in *YM* in Coll-MA based structures can be attributed to the heterogeneity in the material distribution after printing in comparison with BisSR/CEA (see right panel in Figure 2c). Additionally, we determined a compression of the gel-like shell of Coll-MA structures by a factor of ∼4, as the height profile of the structure was significantly influenced by the applied force (see **Figure S4**).

**Figure 2.**
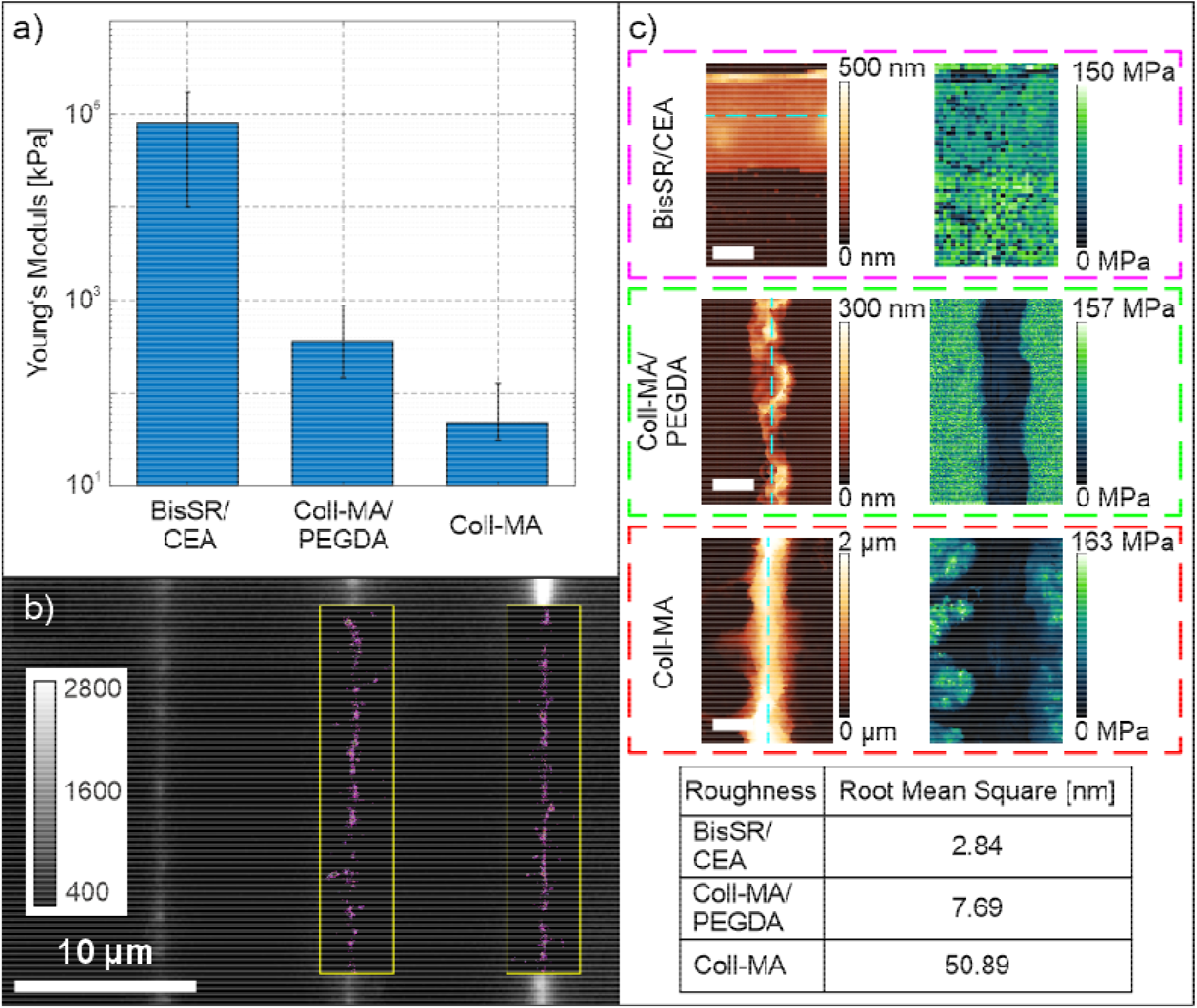
Multi-modal characterization of collagen-based MPL-structures. a) BisSR/CEA *YM* was measured to be 81 ± 10 MPa, 368 ± 150 kPa for Coll-MA/PEGDA, and 49 ± 32 kPa for pure Coll-MA. b) Fluorescence image of suspended Coll-MA lines. The Coll-MA has been labeled with anti-collagen type I IgG Alexa647. SMLM was used to detect single emitters showing a homogenous distribution of collagen (estimated 49.4 ± 11.2 emitters µm^-1^, *N*_*samples*_ = 9) and proved the functionality of collagen after printing. The yellow inset shows a super-resolved image of the lines, with a line width (*i*.*e*. full width at half maximum) of 243 ± 34 nm. c) AFM height and nanoindentation images of BisSR/CEA, Coll-MA/PEGDA and Coll-MA. The table shows the Root Mean Square (RMS) roughness of BisSR/CEA, Coll-MA and Coll-MA/PEGDA along the white dashed lines. The RMS roughness was measured at 2.84 nm for BisSR/CEA, 7.69 nm for Coll-MA/PEGDA and 50.89 nm for Coll-MA. The Coll-MA line exhibits filaments spreading from the core of the line. Scale bar 2 µm for BisSR/CEA and 1 µm for Coll-MA and Coll-MA/PEGDA.

The bioactivity of scaffolds in tissue engineering is as crucial as their mechanical properties. To evaluate the biofunctionality of the Coll-MA lines after printing, we used anti-collagen type 1 IgG Alexa647 for immunolabeling of the lines (Figure 2b). Homogenous spreading of bound antibodies along the written lines was observed *via* SMLM, with an estimated antibody density of 49.4 ± 11.2 emitters µm^-1^ (*N*_*samples*_ = 9) (yellow insets in Figure 2b). The single emitters were localized with a lateral position accuracy of 44 ± 12 nm (17.32 signal-to-noise ratio). Additionally, we estimated the width of lines (at full width at half maximum) to be 243 ± 34 nm, proving sub-diffraction limited feature sizes for Coll-MA based structures for the first time. (for comparison to results from the AFM measurements see Figure S4). Despite the homogenous spreading of antibodies, we found that the Root Mean Square roughness of pure Coll-MA differs from Coll-MA/PEGDA lines by a factor of ∼6, highlighting the homogeneity in the material in contrast with a homogenous surface (see Figure 2c).

To show the effect of a biodegradation-driven change in geometry on cell expansion during 21 days of cell culture, we tested the biodegradation of Coll-MA in hybrid cell cages. The difference in RMS roughness did not influence the time scale of cells degrading the Coll-MA and Coll-MA/PEGDA structures (for results 11 days post-seeding see **Figure S5**a). Once the cells degraded the collagen (typically 6 days post-seeding), further expansion throughout the scaffold was observed (see Figure S5b). While no significant difference for Coll-MA and Coll-MA/PEGDA in biodegradation was observed in a period of 11 days, the surface roughness variations may influence early cell adhesion and degradation, affecting the cellular response. In contrast, cells within the hard cell cages experience lateral spreading restrictions for the entire cultivation period, permitting only axial growth. We show the use of Coll-MA for the fabrication of 3D scaffolds with a biodegradation-driven change in geometry and tunable mechanical properties, effectively mimicking the characteristics of various tissue types.

### 2.3. Vinculin Distribution as an Indicator for Cell-Matrix interactions

Next, we focused on the comparison of cell-matrix interactions in hard collagen-coated BisSR/CEA and hybrid BisSR/CEA in combination with Coll-MA cell cages. To investigate the effect of scaffold stiffness and geometry on cell-matrix interactions, we examined the actin cytoskeleton and the vinculin distributions in MSC. For that, we performed confocal and SMLM imaging of stained MSC 6 and 11 days post-seeding (for confocal images see **Figure 3**a; for SMLM images see **Figure S6**). Our results on hard-/hybrid cell cages verified proper cell adhesion, as the actin filaments associated and co-localized with the vinculin clusters formed at the substrate 6 days post-seeding. Furthermore, we used SMLM data of vinculin for quantitative analysis of mechanosensing at the molecular level. Vinculin was stained using fluorescently labeled (Alexa647 or Alexa488) anti-vinculin IgG. Figure 3b depicts exemplary images of the vinculin distribution in cells within hybrid cell cages 6 days post-seeding (right panel) and within hard cell cages 11 days post-seeding (left panel). The single IgG conjugated Alexa647 emitters in samples 6 days post-seeding were localized with an average position accuracy of 45±9 nm lateral and 109±22 nm axial (signal-to-noise ratio 13.62) and the single IgG conjugated Alexa488 emitters in the samples 11 days post-seeding with an average position accuracy of 49±9 nm lateral and 83±15 nm axial (signal-to-noise ratio 17.02), respectively.^[96]^ As shown in Figure 3c, the vinculin spatial distributions have been arranged into clusters,^[97]^ showing the estimated molecular organization of vinculin within a single cell (**Figure S7** for exemplary clusters). We observed that, for some cells at an early stage, the z-position of cluster centroids (*z*_*pos*_) indicates the presence of two distinct populations, an upper-(yellow dots in Figure 3d) and a lower-one (blue dots in Figure 3d). The upper population primarily comprises protein clusters involved in cell-scaffold interactions, while the lower one contributes to cell adhesion on the glass substrate (Figure 3d). 11 days post-seeding, the upper population redistributes towards the top of the cell (for population filtering see **Figure S8**). As shown in Figure 3e, the average *z*_*pos*_ for all clusters was at 153 ±99 nm above the substrate for 6 days post-seeding and 510 ±154 nm for 11 days post-seeding (for further data of *z*_*pos*_ see **Figure S9**). No correlation between the scaffold version (*i*.*e*. hard/hybrid cell cages) and *z*_*pos*_ was observed at this early developmental stage. The 2CALM-based cluster comparison performed at the single-molecule level,^[98]^ revealed that all clusters are similar in density distribution, regardless of cell cage composition and *z*_*pos*_ (for 6 and 11 days post-seeding). Overall, a 73% similarity of cluster density distribution between all analyzed samples has been determined (Figure 3f). An additional single-molecule cluster analysis comparing the upper-population (6 days post-seeding) and the overall vinculin population (11 days post-seeding), revealed no significant differences (see **Figure S10**). Our results showed gradual increments of *z*_*pos*_, suggesting a similar trend in upper cell layers. However, methodological limitations (such as low signal-to-noise ratios and z-range restrictions) hindered the acquisition of SMLM data regarding the vinculin distribution above 1 µm of the substrate. Additional non-quantitative confocal imaging showed a reduced vinculin population within the cell redirected from the substrate towards the top of the scaffold, along the growth direction (see **Figure S11**). Moreover, confocal images revealed that cells, forming the uppermost cell layer, exhibit a vinculin distribution directed towards the cells beneath them, instead of direct interaction with the scaffold (see **Figure S12**). This finding indicates the unsynchronized growth and development of cells within 3D scaffolds.

**Figure 3.**
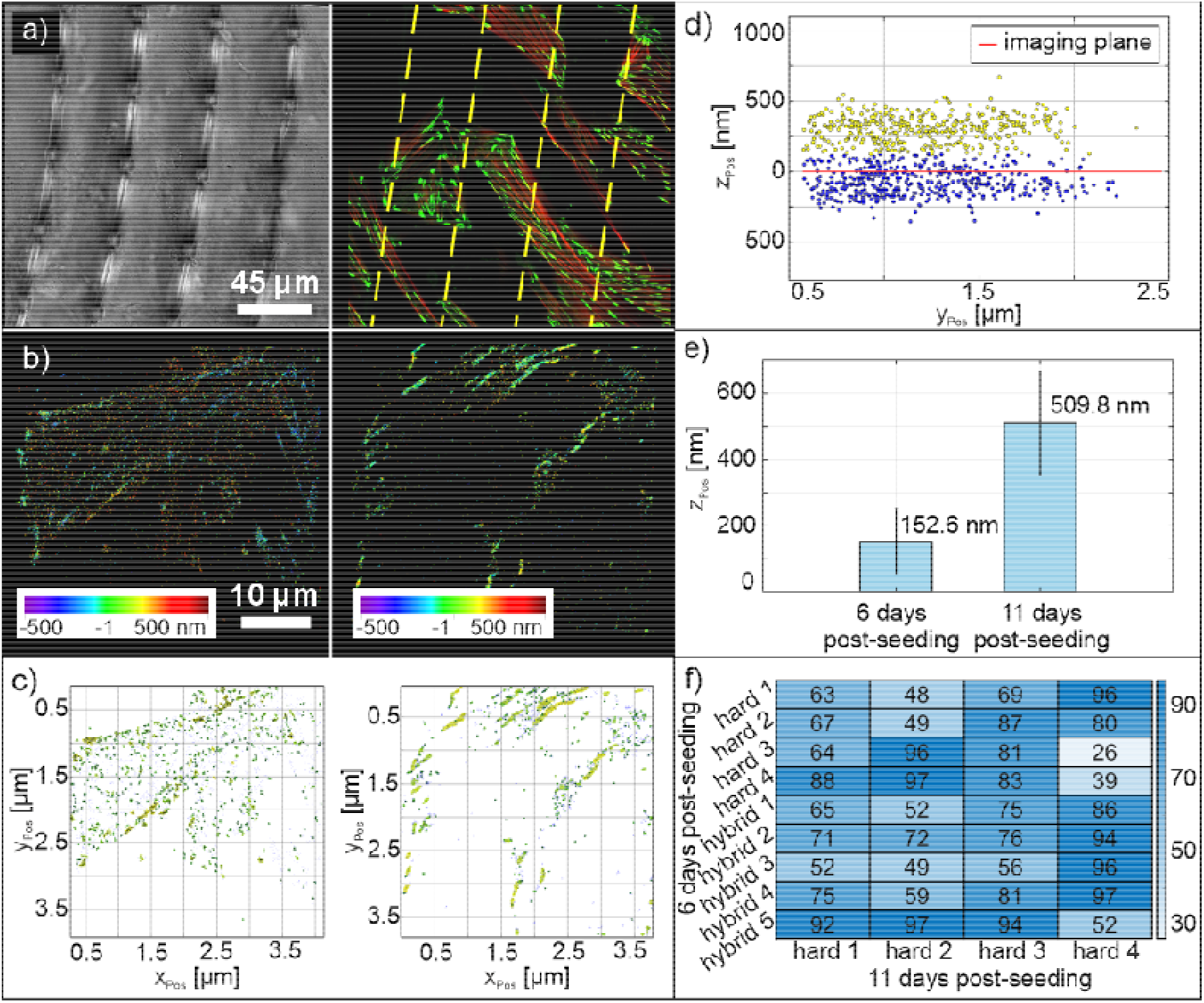
Vinculin cluster formation in 3D scaffolds over time. a) White light image and the corresponding confocal image of vinculin (green) and the actin cytoskeleton (red) of MSC in hybrid cell cages 6 days post-seeding, showing a healthy cell state within the cell cages.^[99]^ b) 3D dSTORM images showing the distributions of vinculin in MSC cultivated in cell cages for 6 days (left) and 11 days (right). Vinculin was labeled with anti-vinculin IgG Alexa647 in samples 6 days post-seeding and anti-vinculin IgG Alexa488 in samples 11 days post-seeding, respectively. The focus was set 1 µm above the substrate. c) The image shows vinculin clusters identified using DBSCAN clustering algorithms (with at least 5 nearest neighbors) in hybrid (6 days post-seeding, left) and hard (11 days post-seeding, right) cell cages.^[97]^ The localized emitter signals are clustered with a radius of *r* = 230 nm (left) and 100 nm (right). d) A distinct vinculin cluster sub-population above the substrate indicates the axial migration of vinculin clusters in MSC 6 days post-seeding over time. e) The average *z*_*pos*_ was 153 ±99 nm in 6 days post-seeding samples, while in 11 days post-seeding samples, *z*_*pos*_ was 510 ±154 nm, indicating a gradual redistribution (*N*_*samples*_ = 9). f) A matrix showing the similarity (in %) of vinculin cluster density distribution for cells 6 days post-seeding and 11 days post-seeding analyzed using 2CALM.^[98]^ The results indicate an overall 73% similarity in vinculin density distributions (*i*.*e*. cluster density and curvature), regardless of the cell cage composition. The plot was generated for DBSCAN cluster radii ranging from 10 to 1000 nm in 30 nm increments.

The cluster analysis showed two relevant results: Firstly, the absolute nm-dimension of the clusters, to investigate the role of vinculin in mechanosensing and the association with the actin cytoskeleton at adhesion sites. Secondly, the vinculin cluster comparison for 6- and 11-days post-seeding in hard and hybrid cell cages showed that there is no significant difference between vinculin clusters in hard and hybrid cell cages; however, we found that *z*_*pos*_ of the vinculin clusters increases over time.

### 2.4. Effect of 3D scaffolds on osteogenic differentiation of MSC

Herein, we address the role of mechanosensing as a response to ECM cues, which are crucial for the development and differentiation of stem cells. Osteogenic differentiation is well characterized and depends on a precisely controlled environment, including medium composition, spatial constraints, and ECM stiffness.^[33,70,79,90]^ However, there is currently a lack of studies and tools, tracking the underlying mechanisms at the single-cell level. Building on our previous results, where we showcased the redistribution of vinculin clusters over time (see chapter 2.3), here our aim is to demonstrate the potential of the 3D scaffold for studying osteogenesis and its link to mechanotransduction. We visualized the formation of vinculin clusters to determine whether vinculin organization influences MSC differentiation and to further investigate the potential impact on osteogenesis. Therefore, we tested the effect of our scaffolds on the expression of osteogenic markers: ALkaline Phosphate (ALP), a marker for confirmation of early osteogenic commitment; collagen type I, an early matrix protein, involved in the early stage of bone formation; osteocalcin, a late-stage marker indicating bone maturation. Firstly, we seeded MSC onto 3D hard cell cages and differentiated them using osteogenic medium. Interestingly, already after 14 days of osteogenic treatment, we saw an increased ALP activity within the cell cages compared to the cells growing in a 2D environment and to cells without osteogenic treatment (See **Figure S13**). Secondly, to compare the results of osteogenic differentiation with our previous findings on cell-matrix interactions within 3D hybrid scaffolds over time, we performed immunostaining of MSC after 21 days under proliferative or osteogenic conditions. Vinculin was stained as outlined above. Collagen type 1 was labeled using either anti-collagen type 1 IgG Fluor CoraLite Plus488, or anti-collagen type 1 IgG Alexa488. Osteocalcin was visualized using primary IgG rabbit anti-osteocalcin and secondary IgG goat anti-rabbit Alexa647. Confocal imaging of vinculin localization in cells treated with expansion medium and osteogenic medium for 21 days revealed no difference regardless of treatment (see Figure S12). However, cells treated with an osteogenic medium exhibited strong expression of both markers (**Figure 4**a). In contrast, cells treated with expansion medium expressed less collagen type 1 and osteocalcin expression was absent (see **Figure S14**). Müller *et. al*. ^[100]^ demonstrated that stem cell repopulation is most effective in single-cell compartments in the short term (7 days), highlighting spatial restrictions as a simple yet powerful tool for directing stem cell fate. Our results showed decreased cell spreading in the hard cell cages (70% of cell cages occupied) under osteogenic conditions compared to cells under expansion conditions and only a subset of the cells (30% of the filled cell cages) expressed both markers (Figure 4b). Yet, our observation of osteogenic marker expression at the border of the 3D scaffold consisting solely of hard cell cages, suggests that spatial constraints influence stem cell differentiation, aligning with Müller *et al*.’s findings. The combination of hard cell cages (yellow inset in Figure 4) and hybrid cell cages (red inset in Figure 4) allowed us to study the impact of a biodegradation-driven change of geometry on the cell spreading within the same scaffold. In the hybrid cell cages, after biodegradation of the Coll-MA structures, cell expansion along the axis of the hard bars of the scaffold was observed (Figure 4c). Collagen type 1 expression was increased by a factor of ∼2 in the hybrid cell cages (average intensity over the scaffold area = 54.61 ± 6.83 a.u. pixel^-1^) compared to the hard cell cages (average intensity over the scaffold area = 23.54 ± 6.36 a.u. pixel^-1^). The observed increase in collagen type I signal highlights scaffold-dependent osteogenic trends. Measurements were collected from multiple regions across independent scaffold areas, confirming the consistency of these patterns.

**Figure 4.**
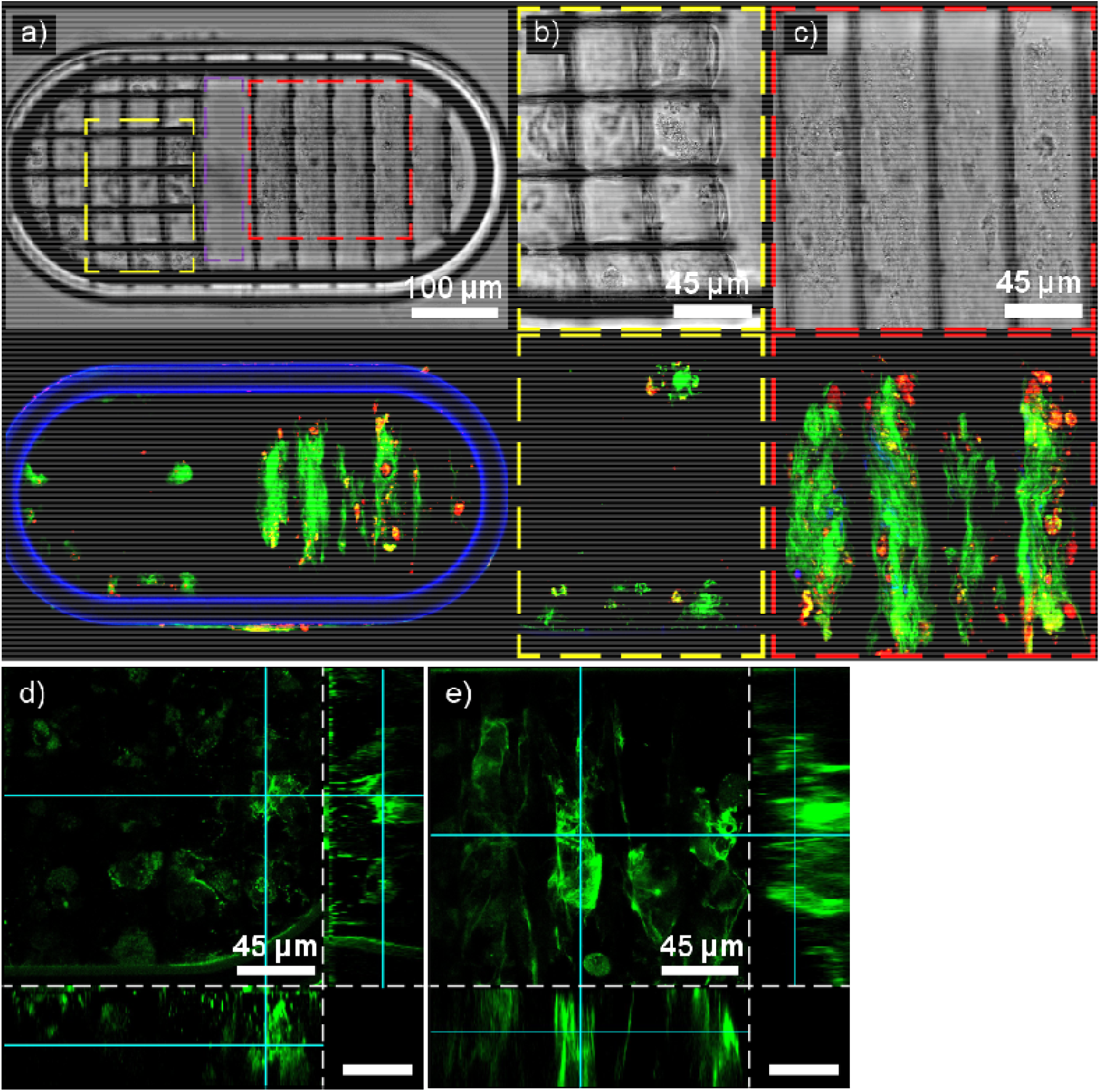
Osteogenic differentiation of MSC in 3D hybrid scaffolds. The upper panel shows the white light images. The lower panel shows the corresponding confocal images. The scale bars hold true for both panels accordingly. ECM was labeled with anti-collagen type 1 IgG Fluor CoraLite Plus488, or anti-collagen type 1 IgG Alexa488 (green), bone mineralization was labeled with primary IgG rabbit anti-osteocalcin and secondary IgG goat anti-rabbit Alexa647 (red), and the cell nuclei were stained with DAPI (blue). Autofluorescence of the stadium-like structure is also shown in blue. a) shows a 3D scaffold with hard and hybrid cell cages. MSC have been in culture and differentiated in osteogenic medium for 21 days. The expansion of MSC and the distribution of collagen type 1 and osteocalcin was scaffold-dependent. Hybrid cell cages showed spreading of the cells throughout the scaffold (red inset), while expansion in hard cell cages was impeded (yellow inset). In the scaffold part with no cell cages, cells were not able to adhere for the 21-day period (purple inset). b) A magnified view on the hard cell cages shows limited cell activity, referring to restricted cell spreading and actin cytoskeleton organization along the scaffold axis, resulting in reduced ECM deposition compared to hybrid cages. Additionally, cells remained spatially constrained with no observable mobility within the hard cages. Only a subset of the cells (30% of cells) within the cell cages formed a collagen type 1 matrix and lower osteocalcin expression was observed compared to MSC in hybrid cell cages. c) In the hybrid cell cages, the cells biodegraded the Coll-MA bars, allowing the cells to expand along the remaining hard part of the cell cage. The hard BisSR/CEA structure supported cell spreading in the z-direction. Distinct clusters of osteocalcin have been detected throughout the hybrid cell cages, indicating successful osteogenic differentiation. d) Confocal image of collagen type 1 in the hard cell cages. The impeded differentiation of the MSC resulted in expression of collagen pre-dominantly near the substrate e) Confocal image of collagen type 1 in the hybrid cell cages. After biodegrading the collagen, the cells were able to expand across the whole scaffolds resulting in collagen expression across the entire scaffold height. Z-axis in (xz)- and (yz)-scans scaled for better readability, scale bar 20 µm.

Analyzing the cell expansion in the z-direction, we found that cells in the hard cell cages expressed collagen type 1 predominantly near the substrate (Figure 4d). In hybrid cell cages, we observed collagen type 1 and osteocalcin expression over the entire scaffold height (Figure 4e). Our previous results showed time- and spatial independence in vinculin clustering of MSC (see chapter 2.3). However, osteogenic markers in hard and hybrid cell cages revealed distinct patterns, revealing a strong dependence on scaffold properties. Despite the importance of ECM stiffness and the mechanotransduction of cells during osteogenesis,^[79]^ the lack of a clear correlation between the vinculin distribution and osteogenic markers highlights the complexity of osteogenesis.^[101–103]^

We observed a correlation in the shape and spreading of cells in hybrid scaffolds, and increased marker expression was determined. Our results demonstrate that the combination of hard (as supporting structure) and biodegradable soft materials (guiding cell expansion) facilitate guided cell expansion over time and thus enabling the fabrication of 3D hybrid scaffolds for tissue growth in any arbitrary 3D geometry.

## 3. Conclusion

We present a modular 3D tissue scaffold with biodegradation-driven change in geometry for OoC applications, designed to investigate cell-matrix interactions at the single-cell and single-molecule levels during expansion and osteogenic differentiation. This modularity provides a novel platform to link single cell mechanosensing to downstream differentiation outcomes in a physiologically relevant 3D context. Our scaffolds integrate biocompatible synthetic polymer (BisSR/CEA) with bioactive Coll-MA, enabling stiffness tuning from ∼50 kPa to over 80 MPa, while providing mechanical and biochemical cues that promote stem cell differentiation towards the osteogenic lineage. Using multi-modal analysis, we systematically characterized nanoscale Coll-MA structures. We demonstrate the tunability of the *YM* through PEGDA incorporation. Furthermore, the biodegradable nature of Coll-MA enables the design of 3D scaffolds with biodegradation-driven change of geometry, facilitating guided cell expansion. We show the gradual migration of vinculin clusters in MSC over time. The similarity of clusters observed in both hard and hybrid microenvironments suggests a generalized adaptive response rather than stiffness-specific mechanotransduction, indicating a predominant influence from the rigid glass substrate. Notably, these findings were achieved using single-molecule sensitive techniques, offering a more detailed understanding of vinculin dynamics and cellular behavior compared to previous studies. In contrast to findings by Lo *et. al*.^[104]^, reporting a strong association between increased vinculin clustering and accelerated osteogenesis in single cells, our results revealed only a weak correlation between vinculin clustering and the expression of osteogenic markers. This highlights the complexity of osteogenesis, which is influenced by mechanical and biochemical cues, such as ECM composition, cell density, scaffold architecture, and timing.^[105]^

We observed that MSC in hybrid scaffolds exhibit enhanced osteogenic marker expression, while in the hard scaffolds, a second population of non-osteogenic cells was discovered. This emphasizes the importance of modular 3D environments in guiding stem cell fate beyond chemical cues, providing insights that may not have been captured in earlier research. Importantly, in contrast to many prior 3D mechanobiology studies that relied on bulk or multicellular averages^[75–78]^, our results show that local geometry and bioactivity—rather than overall scaffold stiffness—directly regulate osteogenic differentiation at the single-cell level. Although current research was still limited in geometry, size, limited biofunctional components, and multi-cell-type support challenges, the results highlight the potential of modular scaffolds for studying the environment-dependent development (or differentiation) of various cell phenotypes.

The scaffold does not replicate the full hierarchical structure of bone, such as osteocyte lacunae or canaliculi, but provides controlled single-cell environments for studying mechanosensing, with potential for integration of more complex microarchitectures in the future.

Future developments on expanding co-culture capabilities, incorporating additional bioactive cues, and integrating with dynamic microfluidic systems can offer a promising strategy for developing complex tissue interfaces, *e*.*g*. the bone-cartilage junction. The platform can potentially address the correlation of mechanical and biochemical cues behind tissue development (*e*.*g*., osteogenesis). Further molecular-level studies can enhance understanding of tailored microenvironments, driving advancements in tissue engineering and personalized regenerative medicine for bone and cartilage repair, support bone-implant integration, and facilitate disease modeling for conditions like osteoporosis and osteoarthritis.

Looking forward, MPL-based modular scaffolds could enable experiments that remain inaccessible to conventional methods, for example: (i) single-cell level mechanosensing in architecturally complex niches, where local degradability and rigidity can be varied independently, (ii) time-resolved studies of cell fate in dynamically remodeling microenvironments, with degradation-induced geometry changes controlled by design rather than random hydrogel breakdown, and (iii) design of graded or anisotropic mechanical environments that mimic native bone–cartilage or tendon–bone interfaces, with single-cell resolution analysis of differentiation along the gradient. These unique capabilities highlight the advantage of MPL-based scaffold design over traditional hydrogel or bulk scaffold approaches and underline its potential to open previously inaccessible mechanobiological experiments.

## 4. Experimental Section/Methods

### Materials

Ormocomp® (Micro Resist Technology) was used as monomer for the stadium-like structure. For the hard cell cages and the hard part of the hybrid cell cages, a mixture of the monomers of Bisphenol A-Glycidyl MethAcrylate (Bis-GMA, Esschem Europe), ethoxylated bisphenol A dimethacrylate (SR348C, SARTOMER) and 2-CarboxyEthyl Acrylate (CEA, Sigma Aldrich) in a weight ratio of 1:3.5:0.1 were used (BisSR/CEA). 1 wt% of Irgacure 819 (Sigma Aldrich) was added as photoinitator.

For the soft part of the hybrid cell cages, methacrylated collagen (Coll-MA, Sigma Aldrich) was dissolved at a concentration of 6 mg mL^-1^ in 20 mM acetic acid over night at 4 °C. 0.25 wt% Riboflavin (Vitamin B_2_) (Sigma Aldrich) was added prior to printing as photoinitiator. 5 wt% of Poly(Ethylene Gycol) DiAcrylate (PEGDA, M_n_ = 575, Sigma Aldrich) was added as a crosslinking agent for Coll-MA/PEGDA structures.

### Multi-Photon Lithography (MPL)

MPL was performed with a customized lithography system (Workshop Of Photonics (WOP)) using an ultra-short pulsed laser at 515 nm (CARBIDE, 1 MHz repetition rate, >290 fs pulse duration, Light Conversion) and a 3-axis stage (AEROTECH Nanopositioner) for sample movement, as described previously.^[19]^ Three-dimensional MPL scaffolds were prepared using an air objective lens (50x, 0.42 NA, Mitutoyo). The stadium-like structure and the supporting bars for the suspended lines were fabricated with an excitation intensity of 0.067 TW cm^-2^ and a writing speed of 1 mm s^-1^, followed by rinsing with 100% acetone (Roth). BisSR/CEA structures were printed as follows: pillars were written with an excitation intensity of 0.038 TW cm^-2^ at 1 mm s^-1^ writing speed while bars were generated via double illumination of each voxel with an excitation intensity of 0.059 TW cm^-2^ at the same writing speed. BisSR/CEA structures were developed by rinsing with 100% ethanol (Roth). Coll-MA/PEGDA structures were fabricated using double illumination of each voxel with an excitation intensity of 0.084 TW cm^-2^ and 0.1 mm s^-1^ writing speed. Similarly, Coll-MA structures were produced with a double illumination of each voxel with an excitation intensity of 0.091 TW cm^-2^ and a writing speed of 0.1 mm s^-1^. Coll-MA based structures were developed by gentle washing with 0.5 mM acetic acid (Sigma Aldrich) and stored in ultra-pure water until further use.

### MSC from Wharton’s jelly culture

MSC were thawed and expanded in MSC expansion medium (StemMACS, Milteny Biotec) supplemented with MSC-CytoMix (Milteny Biotec) and 1% penicillin-streptomycin (P/S). MSC were passaged using 0.25% Trypsin Versene (VWR) and centrifuged at 220xg for 5 min. The cells were expanded twice before seeding on 3D scaffolds.

### MSC seeding on 3D scaffolds

Prior to cell seeding, the samples were sterilized with UV-light for 30 min, washed 3x times with sterile phosphate buffered saline without Ca/Mg (PBS, Roth). 3D scaffolds with only hard cell cages were coated with 50 µg mL^-1^ collagen type 1 from rat tail (Ibidi) in 17.5 mM acetic acid for 15 min prior to seeding. 10 000 cells were seeded on the scaffolds in MSC expansion medium and kept at 37 °C and 5% CO_2_ for 6 to 11 days, exchanging the medium at least every 2 days.

### Osteogenic differentiation of MSC

For osteogenic differentiation, MSC were seeded at a density of 4 000 cells per scaffold in MSC expansion medium (StemMACS, Milteny Biotec) and incubated overnight. Next day, medium was exchanged to osteogenic differentiation medium DMEM-low glucose (Thermo Fisher) containing 10% fetal calf serum (FCS, Sigma Aldrich), 2 mM L-glutamine (Thermo Fisher), 100 U mL^-1^ P/S (Thermo Fisher), 10 nM dexamethasone (Sigma Aldrich), 150 μM ascorbat-2-phosphate (Sigma-Aldrich), 10 mM β-glycerophosphate (StemCell Technologies) and 10 nM dihydroxy-vitamin D3 (Sigma-Aldrich) or control medium consisting of DMEM-low glucose, L-glutamine with 10% FCS and 100 U mL^-1^ P/S. Medium was changed every 3-4 days.

### Immunostaining

MSC were washed with PBS at 37 °C 3 times, fixed using 4% paraformaldehyde (Arcos Organics) in PBS at 37 °C for 20 min, permeabilized in 0.5% Triton X-100 in PBS and blocked in 1% albumin from chicken egg white (ACE, Sigma-Aldrich) for 1 h. The actin cytoskeleton was labeled with 6.6 nM phalloidin Alexa647plus (Thermo Fisher) in ACE for 20 min. Vinculin was labelled either with 1 µg mL^-1^ anti-vinculin IgG Alexa488 (Abcam) or 1 µg mL^-1^ anti-vinculin IgG Alexa647 (Abcam) in ACE for 1 h. Collagen type 1 was labelled with 5 µg mL^-1^ anti-collagen type 1 IgG Fluor CoraLite Plus 488 (Fisher Scientific) or 1 µg mL^-1^ anti-collagen type 1 IgG Alexa647 (Abcam) in ACE for 1 h. Osteocalcin was labelled with 10 µg mL^-1^ primary IgG rabbit anti-osteocalcin (Abcam) in 1% ACE overnight at 4 °C and 2 µg mL^-1^ secondary IgG goat anti-rabbit Alexa647 (Fisher Scientific) in ACE for 1 h. The cell’s nuclei were stained with 2 µg mL^-1^ DAPI (Thermo Fisher) in PBS for 15 min.

### Fluorescence microscopy

Fluorescence images were acquired using a modified Olympus IX81. Precise sample positioning was done using a XYZ piezo stage with a nanometer precision (Physik Instrumente GmbH) in combination with a mechanical stage with a 1x1 cm range (JPK Instruments) for coarse movement. MSC were illuminated through a 60x magnification objective lens (1.4 NA, Olympus) and an additional 1.6x magnification tube lens for a final magnification of 96. For 3D imaging, a cylindrical lens (f = 1 000 mm, Thorlabs) was used in the optical detection pathway of the microscope. A 642 nm solid-state diode pumped laser (Toptica Photonics), a 488 nm solid-state pumped diode laser (Toptica Photonics), and a 405 nm diode laser (Insaneware) were used as light sources. A dichroic filter (ZT405/488/561/640rpc nm, Chroma) and an emission filter (446/523/600/677 nm BrightLine quad-band band-pass filter, Semrock) were used. An additional emission filter (ET 700/75 M, Chroma Technology GmbH) was used for imaging of phalloidin Alexa647plus, anti-vinculin IgG Alexa647 and an additional emission filter (ET 525/50 M, Chroma Technology GmbH) was used for imaging of anti-vinculin IgG Alexa488. Fluorescence signals were collected with an Andor iXonEM+ 897 (back-illuminated) EMCCD camera (16 μm pixel size).

For 3D imaging with nanometer accuracy, direct STochastic Optical Reconstruction Microscopy (dSTORM) with a cylindrical lens in the optical detection path was implored. Calibration of the axial-dependent deformation of the point spread function was done by using a sample of homogeneously distributed TetraSpeck™ beads (0.1 µm, Invitrogen) on a glass cover slip. 200 images were taken at a defined step size of 10 nm along the axial axis. 10 000 images were acquired per imaging sequence with 20 ms illumination time and a sampling rate of 25 images s^-1^. The single-molecule signals were analyzed and visualized using a custom written software.^[96]^

Confocal imaging was performed with a Leica Stellaris microscope using either an air objective lens (20x, 0.75 NA), or a glycerin immersion objective lens (63x, 1.30 NA). The 3D stacks were recorded by illuminating the sample with 400 nm, 491 nm, 640 nm laser light. Images were analyzed with Fiji/ImageJ V2.

### Cluster analysis

For comparison of the 3D vinculin localization data, clustering-based analysis using 2CALM was performed.^[98]^ The 2-sample Comparative Analysis of 3D Localization Microscopy Data platform (2CALM) is an analysis pipeline that organizes localization microscopy data into protein clusters of varying dimensions and calculates statistical parameters using various numerical methods. Localization microscopy images were treated as 3D point clouds, and a comparative analysis required extracting features that represent their geometric structure without compromising accuracy. To achieve this, point clouds (vinculin localizations) were clustered within a constant radius. For each cluster, the relative density (compared to the overall density of the sample) and curvature were calculated. These distributions were then pairwise compared using the Kolmogorov-Smirnov and Wilcoxon tests, and the p-values were determined and aggregated. This process generates a curve of p-values for each radius in the given sequence (from 5 nm to 1000 nm, with a step size of 30 nm). Based on this p-value curve, the level of similarity was determined. The platform enabled the assessment of protein (i.e. vinculin) distributions in two biological samples, identifying nanoscale differences through parameters such as cluster density and curvature. Its automated classification system integrated multiplexing and multi-level statistical techniques into a unified parameter, facilitating sample comparison and similarity testing. We employed DBSCAN-based clustering to assess the relationship between density and curvature as a function of cluster size for both samples.^[97]^ The DBSCAN clustering radius was automatically calculated for each sample as the average minimum distance between sample points, multiplied by a factor of 2. This method also enabled the exclusion of noise from non-specifically bound fluorescent antibodies. Furthermore, the cluster centroids were extracted to calculate the relative distance of the clusters above the substrate.

### Atomic Force Microscopy (AFM)

AFM measurements were performed with an JPK Nano Wizard 4 using QI™ mode. The AFM was mounted on top of an inverted epi-fluorescence microscope (AxioObserver, Zeiss). The *YM* of BisSR/CEA and Coll-MA were measured with a PPP NCHR cantilever (Nanosensors, nominal spring constant = 42 N m^-1^) and the *YM* of Coll-MA/PEGDA was measured with a PPP FMR cantilever (Nanosensors, nominal spring constant = 2.8 N m^-1^). The exact cantilever spring constant was determined before each measurement using the contact-based calibration method (JPK).

## Supporting Information

Supporting Information is available from the Wiley Online Library or from the author.

## Acknowledgements

We thank Heidi Piglmayer-Brezina for taking SEM images of the 3D scaffolds. We would also like to thank Dr. Markus Axmann for proofreading the manuscript.

Received: ((will be filled in by the editorial staff))

Revised: ((will be filled in by the editorial staff))

Published online: ((will be filled in by the editorial staff))

