## supplement for "Hybrid MPL Scaffolds with Nanoscale Mechanobiology for Bone-on-Chip"

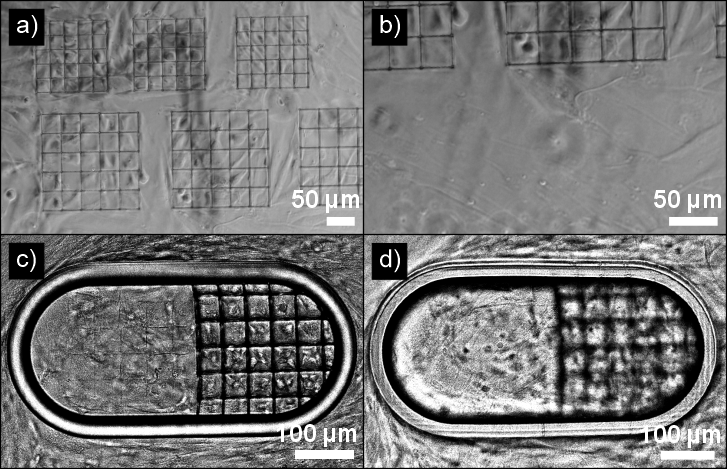

**Figure S1.** Comparison of cell expansion in the absence/presence of the stadium-like structure. a) and b) show MSC growing inside the cell cages and on the glass substrate. In the absence of a barrier, cells inside the cell cages were able to extend beyond the scaffold, possibly migrate through the scaffold and communicate with cells spreading exclusively in a 2D environment. c) and d) show cells growing in cell cages separated by the stadium-like structure during expansion. Imaging at the glass substrate shows the separation of the cells (c) while imaging at the top of the stadium-like structure, shows cells were not able to overgrow the stadium-like structure, effectively isolating the cells in the 3D environment from the cells in the 2D environment (d).

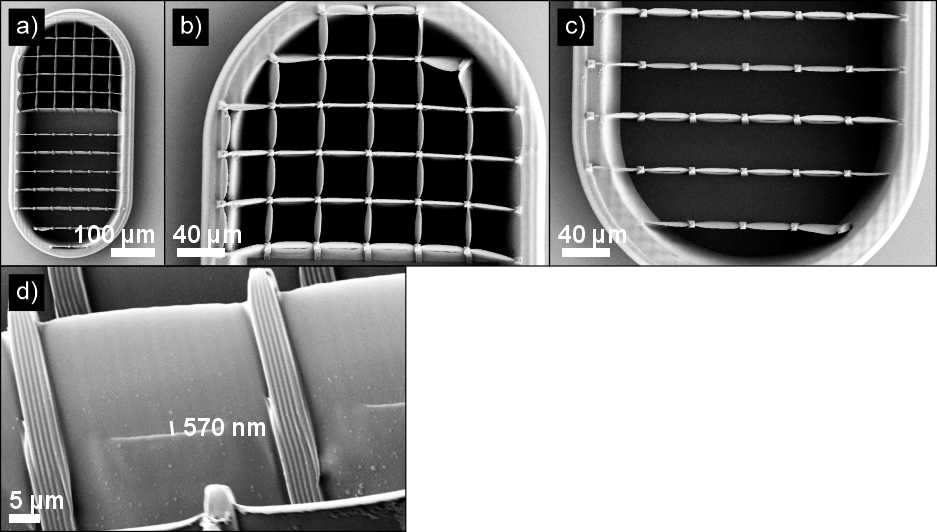

**Figure S2.** SEM images of the 3D hybrid scaffold. a) Overview of the stadium-like structure filled with hard and hybrid cell cages. b) Zoom-in on the hard cell cages. c) Zoom-in on the hard part of the hybrid cell cages. d) Zoom-in on the bars of the hybrid cell cages shows a gap of 570 nm between the bars, allowing cell-cell interactions during cultivation.

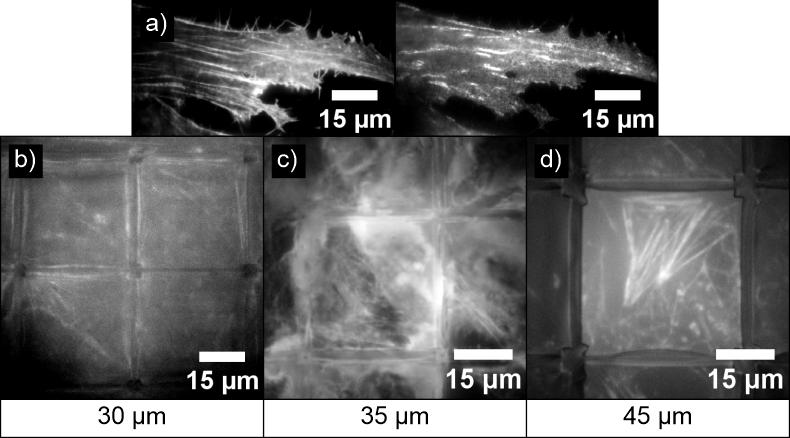

**Figure S3.** Grid size-dependent cell expansion and actin expression in cell cages. Equal scaling of gray values for all images. The actin cytoskeleton of MSC was labelled 6 days post-seeding as an indicator for cell morphology, spreading and cytoskeleton organization. a) shows the actin cytoskeleton and the vinculin of a MSC on a glass substrate as a reference. b) Contrary to the initial spreading of MCS in 30 µm hard cell cages only scarce expression of actin is observed. c) shows the altered actin cytoskeleton of a MSC in a 35 µm hard cell cage. Despite the early expansion of cells, the distorted actin cytoskeleton is a strong indicator for low cell viability.^[1]^ d) shows the actin filaments of a MSC in a 45 µm hard cell cage. The cell expresses well organized actin filaments indicating a healthy cell state.

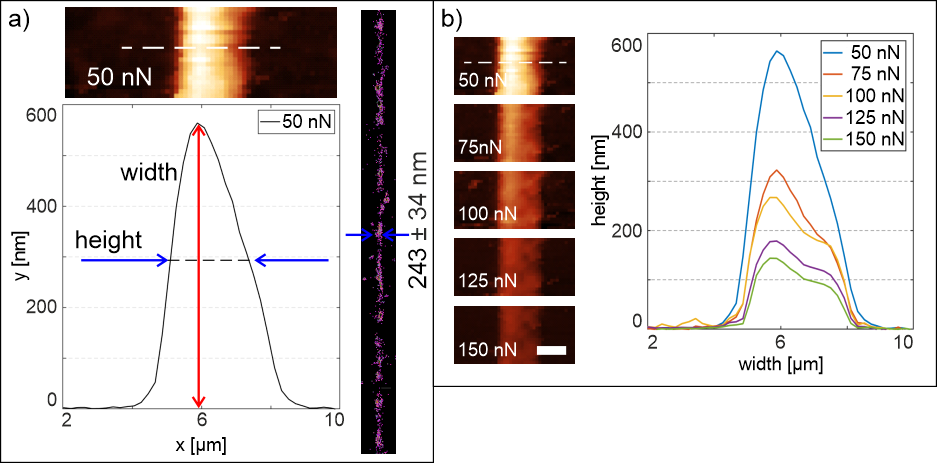

**Figure S4.** Feature size of Coll-MA lines. a) shows the exemplary height profile of a Coll-MA line measured at 50 nN. The measured y-profile of the line with AFM (382 ± 143 nm) is corresponding to the width measured with SMLM (243 ± 34 nm), as the line falls over after printing. Detailed measurement setup can be found in.^[2]^ b) AFM height profile of Coll-MA lines measured with 50 - 150 nN applied force. The measured heights as function of the applied force: 382 ± 143 nm (50 nN), 212 ± 80 nm (75 nN), 190 ± 60 nm (100 nN), 124 ± 44 nm (125 nN) and 96 ± 38 nm 150 nN) were measured. Scale bar 2 µm.

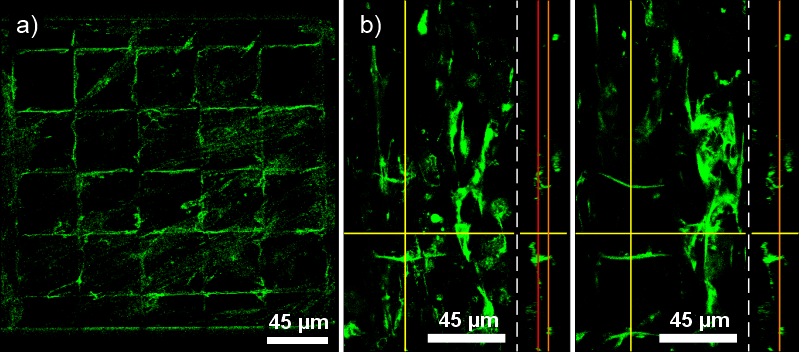

**Figure S5.** Residual Coll-MA/PEGDA cell cages and Coll-MA lines. a) shows the confocal image of the autofluorescence of the remains of Coll-MA/PEGDA cell cages on the glass substrate 11 days post-seeding. The cells metabolized the Coll-MA/PEGDA over that time span. b) shows confocal images of remaining Coll-MA in hybrid cell cages and collagen type 1 in MSC labelled with anti-collagen type 1 IgG Alexa 488 21 days post-seeding. In the lower segment of the cell cages (red line indicates imaging plane), the cells destroyed the Coll-MA lines and were able to expand in the scaffold, while in the higher segment of the scaffold (orange line indicates imaging plane) the Coll-MA lines remained intact, blocking cell expansion in x- and y- direction.

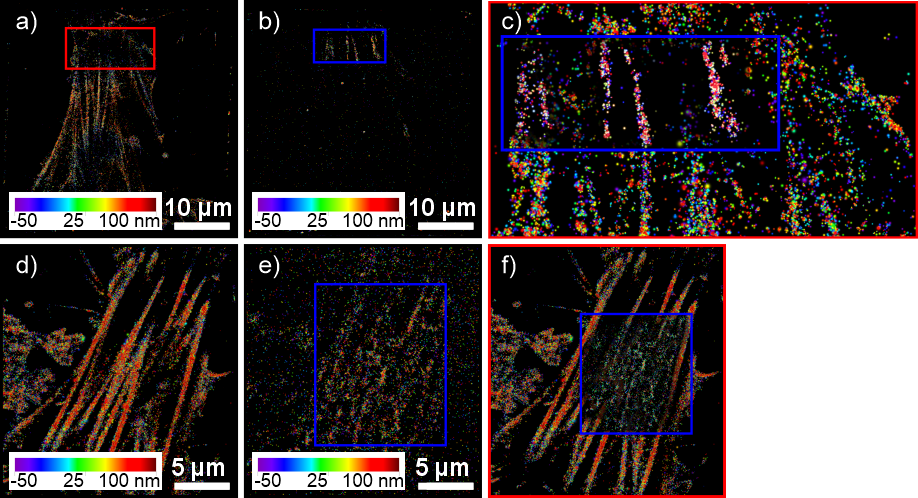

**Figure S6.** Overlay of actin and vinculin images. a,d) 3D STORM image of the actin cytoskeleton of a MSC in a hard cell cage 11 days post-seeding. b,e) 3D STORM image of the vinculin of the same cell. c) Overlay of the insets in a) and b) the vinculin clusters complement the origin of the actin filaments, as the vinculin associates with the actin to regulate the mechanotransduction. f) Overlay of d) and the inset in e) shows the co-localization of the vinculin clusters formed at the substrate with the associated actin filaments.

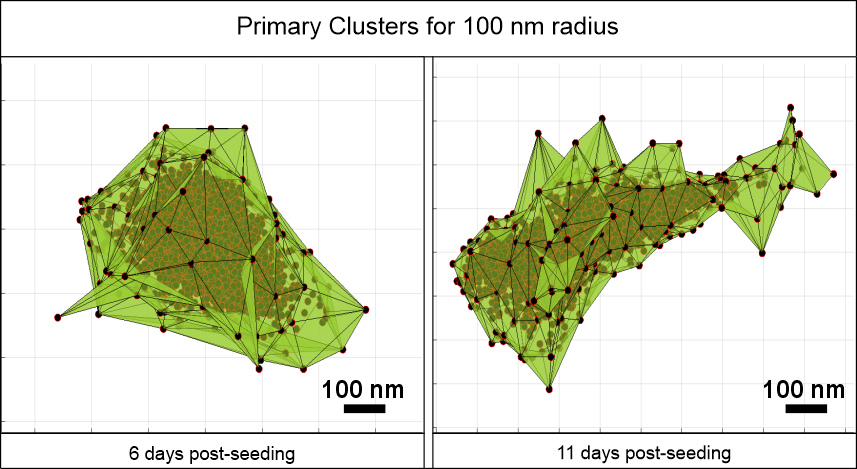

**Figure S7.** Exemplary vinculin clusters of MSC 6 and 11 days post-seeding in hard cell cages for 100 nm clustering radius.

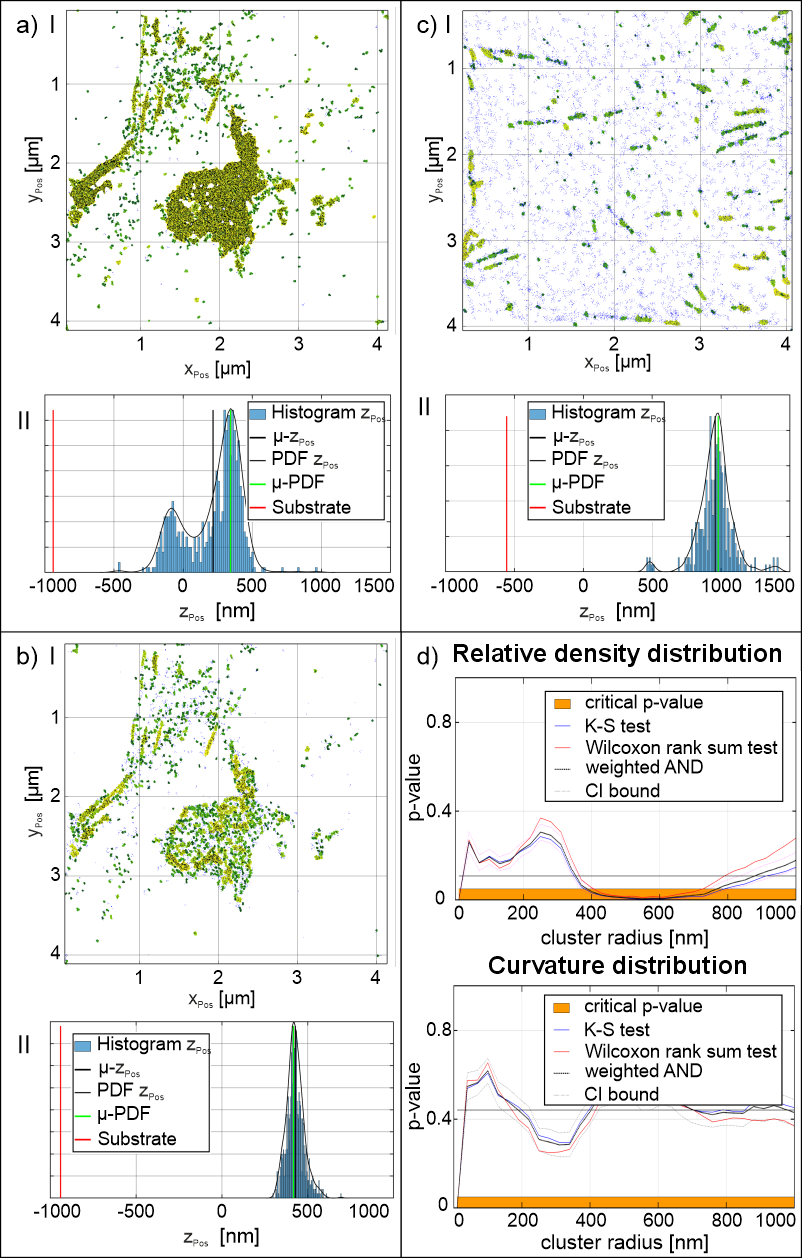

**Figure S8.** Filtering of vinculin clusters and statistical comparison of 6 days post-seeding and 11 days post-seeding in hard cell cages a I) and c I) depict the clustering of vinculin raw data in MSC 6 days post-seeding (clustering radius: 200 nm) and 11 days post-seeding (clustering radius: 100 nm), respectively. The corresponding average distance of the cluster centroids in z-direction (z_pos_) distributions are shown in a II) and c II), revealing two distinct local maxima, which facilitated the classification of vinculin clusters into lower and upper populations. Following this classification, c I) presents the filtered vinculin clusters at 6 days post-seeding, specifically for the upper population (clustering radius: 150 nm), with its corresponding z_pos_ distribution shown in c II). d) provides a statistical comparison of the relative cluster density and curvature distribution for the clusters displayed in b I) and c I). Statistical analyses (Kolmogorow-Smirnov (K-S) test, Wilcoxon rank sum test, weighted AND test of both tests) were performed according to the methodology outlined by Mayr et al.^[3]^ Similarities were as follows: for the relative density distribution: Sim_M_ = 0.77, Sim_L_ = 0.65, average p-value = 0.108 and for the Curvature distribution: Sim_M_ = 0.94, Sim_L_ = 0.97, average p-value = 0.441. The results of these comparisons are presented in Figure 3f and Figure S10.

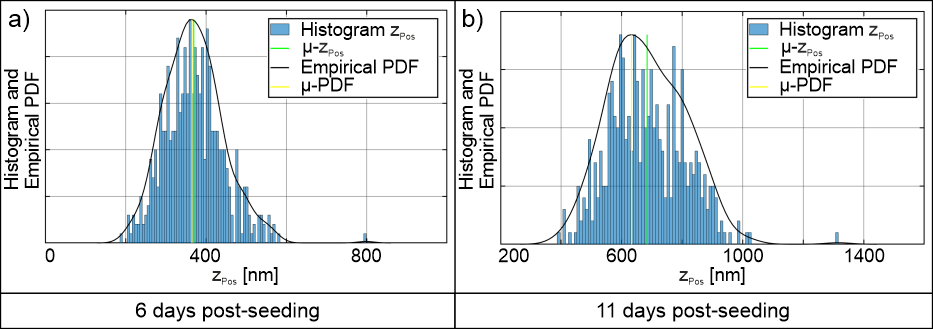

**Figure S9.** Histograms of vinculin cluster centroid distance above the substrate in the z-direction (z_pos_). a) Corresponding z_pos_ distribution of the 6 days post-seeding sample in Figure 3b. The average distance above the substrate of vinculin clusters was at 370 nm while in b) the distance above the substrate for cluster centroids shifted to 684 nm in the 11 days post-seeding sample, indicating the shift of vinculin clusters in the z-direction over time.

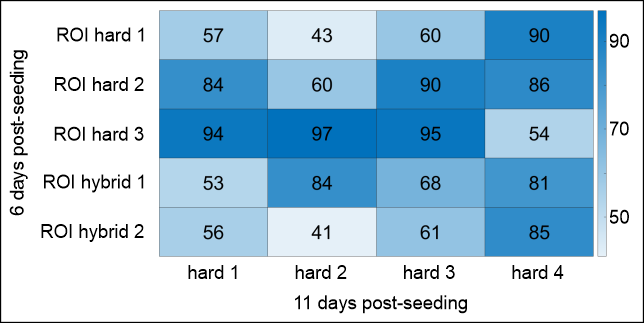

**Figure S10.** 2CALM-comparison of the vinculin clusters upper population in MSC in hard and hybrid cell cages 6 days post-seeding and the vinculin clusters in MSC formed 11 days post-seeding inside MSC in hard cell cages. No significant difference was found between the two populations.

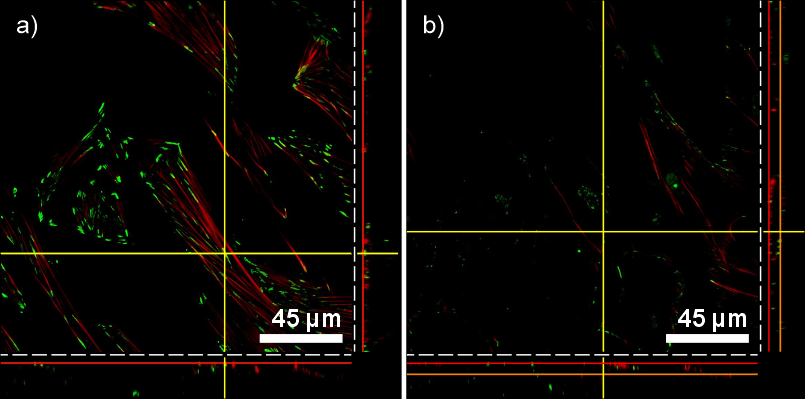

**Figure S11.** Actin and vinculin confocal images in hybrid cell cages. a) shows the distribution of vinculin (green) and actin (red) near the glass substrate (red line indicates imaging plane). Actin filament formation started at the vinculin clusters. b) above the glass substrate (orange line indicates imaging plane) the vinculin cluster density decreases.

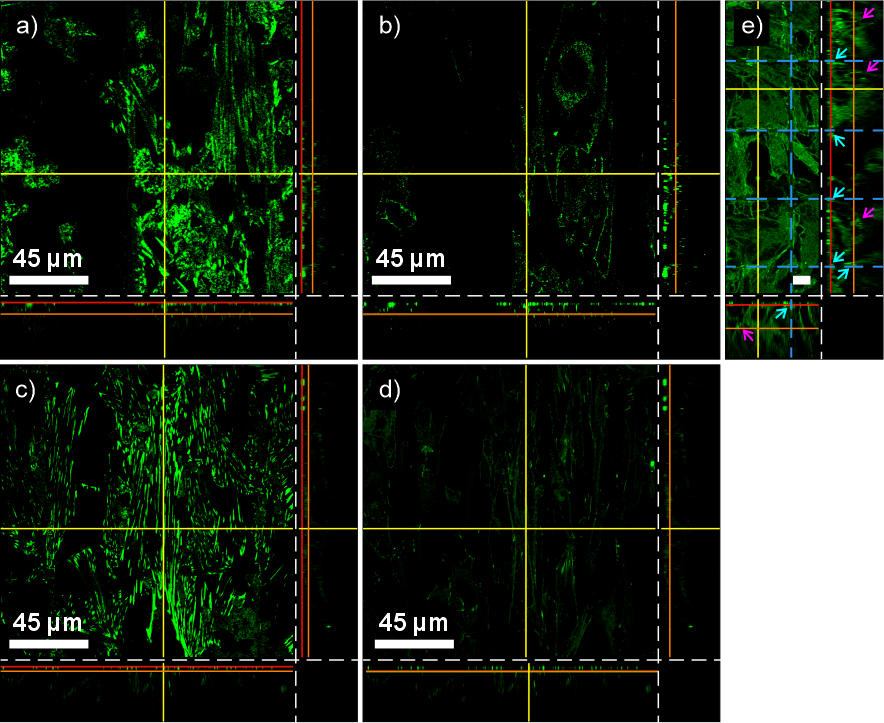

**Figure S12.** Vinculin clusters at the glass substrate and between cell layers during expansion and during differentiation. Red lines indicate first imaging plane/cell layer at the substrate, orange lines indicate second imaging plane/cell layer (optical sectioning z = 4 µm) a, c) shows the confocal image of vinculin labelled with anti-vinculin IgG Alexa488 for cells treated with expansion medium and osteogenic medium, respectively. Vinculin clusters formed on the glass substrate and in regions where the cell interacts with the cell cages. b, d) shows the confocal image at the same x- and y-position at the second cell layer. Most of the vinculin clusters were observed as cell-cell interactions rather than cell-matrix interactions, as seen at the substrate. e) shows a confocal image of cells within hard cell cages treated with expansion medium. Blue dashed lines indicate the scaffold. The arrows highlight cell-scaffold interactions (cyan) and cell-cell interactions (pink), respectively. Scale bar 10 µm.

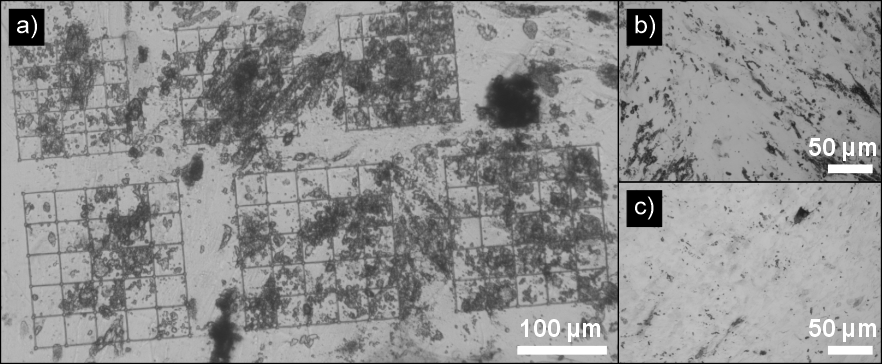

**Figure S13.** ALkaline Phosphatase (ALP) staining of MSC. a) White light image of MSC in hard cell cages after differentiation in osteogenic medium for 14 days. ALP staining was performed with SIGMA FAST BCIP®/NBT protocol. An increase of activity of ALP can observed within the cell cages compared to the surrounding cells with no cell cages. b) MSC solely expanding in a 2D environment showed in increased ALP activity. c) Control experiment with no osteogenic medium.

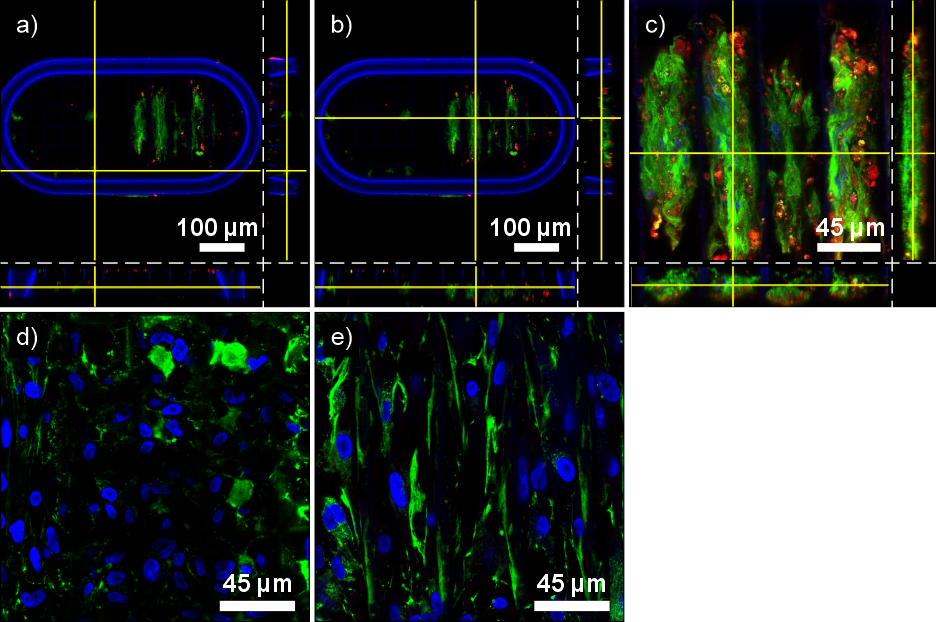

**Figure S14.** Additional confocal images of MSC 21 days post-seeding. a), b) and c) show images of cells treated with osteogenic medium. d) and e) show images of cells treated with expansion medium. ECM, bone mineralization and cell nuclei have been stained with anti-collagen type 1 IgG Fluor CoraLite Plus 488 (green), primary rabbit IgG anti-osteocalcin and secondary IgG goat anti-rabbit Alexa647 (red) and DAPI (blue), respectively. Autofluorescence of the stadium-like structure also in blue in a) and b). a) shows the collagen type 1 matrix within a hard cell cage. The cell expansion is restricted in x- and y- direction. b) shows the collagen type 1 matrix within a hybrid cell cage. The expression of collagen type 1 in the z-direction is higher compared to a). Furthermore, the cells are able to expand across the entire scaffold. c) shows an overlay image of all 3 labels demonstrating the expression of collagen type 1 and osteocalcin across the entire scaffold in z direction. d) and e) show cells treated with expansion medium within hard and hybrid cell cages respectively. Without specific growth factors for osteogenic differentiation, the expression of collagen type 1 is reduced, and the expression of osteocalcin is absent entirely. In comparison with cells treated with osteogenic medium, cells expanded across the whole scaffold and the proliferation is increased.
